# Purifying selection enduringly acts on the sequence evolution of highly expressed proteins in *Escherichia coli*

**DOI:** 10.1101/2022.03.02.482674

**Authors:** Atsushi Shibai, Hazuki Kotani, Natsue Sakata, Chikara Furusawa, Saburo Tsuru

**Author notes:** **Corresponding author:** Saburo Tsuru, Universal Biology Institute, Graduate School of Science, The University of Tokyo, Faculty of Science Bldg.1, Room No.: 446, 7-3-1 Hongo, Bunkyo-ku, Tokyo 113-0033, Japan.

## Abstract

The evolutionary speed of a protein sequence is constrained by its expression level, with highly expressed proteins evolving relatively slowly. This negative correlation between expression levels and evolutionary rates (known as the E–R anticorrelation) has already been widely observed in past macroevolution between species from bacteria to animals. However, it remains unclear whether this seemingly general law also governs recent evolution, including past and *de novo*, within a species. However, the advent of genomic sequencing and high-throughput phenotyping, particularly for bacteria, has revealed fundamental gaps between the two evolutionary processes and has provided empirical data opposing the possible underlying mechanisms which are widely believed. These conflicts raise questions about the generalization of the E–R anticorrelation and the relevance of plausible mechanisms. To explore the ubiquitous impact of expression level on molecular evolution, and to test the relevance of the possible underlying mechanisms, we analyzed the genome sequences of 99 strains of *Escherichia coli* for microevolution in nature. We also analyzed genomic mutations accumulated under laboratory conditions as a model of *de novo* microevolution. Here, we show that the E–R anticorrelation is significant in both past and *de novo* microevolution in *E. coli*. Our data also confirmed ongoing purifying selection acting on highly expressed genes. Ongoing selection included codon-level purifying selection, supporting the relevance of the underlying mechanisms. However, their contributions to the constraints in recent evolution might be smaller than previously expected from past macroevolution.

## Introduction

Is there any general law that governs the evolution of protein sequences on Earth? The rate of protein sequence evolution differs between genes. Many factors other than functional importance have been proposed as determinants for the rate of evolutionary diversification among a protein sequence, as reviewed by Zhang and Yang (2015). Among these factors, gene expression levels might be a general determinant (Krylov *et al*. 2003; Rocha and Danchin 2004; Drummond and Wilke 2008). Comparative genomics of orthologous genes of closely related species revealed a pervasive negative correlation between gene expression level and the rate of evolutionary diversification in a protein sequence, namely E–R (Expression–evolutionary Rate) anticorrelation (Pál *et al*. 2001). The underlying mechanism of the E–R anticorrelation remains unclear (Usmanova *et al*. 2021) but can be explained by several purifying selections, such as the selection against mistranslation and protein misfolding (Akashi 1994; Drummond *et al*. 2005; Drummond and Wilke 2008; Allan Drummond and Wilke 2009; Cherry 2010a; Yang *et al*. 2010; Geiler-Samerotte *et al*. 2011), selection against incorrect and slow translation (Akashi and Gojobori 2002; Cherry 2010b; Gout *et al*. 2010; Park *et al*. 2013; Yang *et al*. 2014), and selection against protein misinteraction (Zhang *et al*. 2008; Levy *et al*. 2012; Yang *et al*. 2012). These purifying selections are believed to be strong for highly expressed proteins because the defects in the quality and quantity of these proteins presumably confer more deleterious effects on the cells than poorly expressed proteins when considering the law of mass action.

Contrary to the ubiquity of the E–R anticorrelation for evolution between species (macroevolution), little is known about whether the same law governs evolution within species (microevolution). Interestingly, the advent of genomic sequencing and high-throughput phenotyping has revealed several gaps between the two evolutionary processes, particularly among bacteria. Notably, bacterial phenotypic diversification in nature is biphasic, whereby phenotypic diversification (such as metabolism) occurs rapidly and instantaneously within species (microevolution), while divergence between species or genera (macroevolution) proceeds gradually (Plata *et al*. 2015). Consistent with this general trend in phenotypes, recent studies have also revealed an unexpectedly large genetic divergence of protein sequences attributable to weaker purifying selection within bacterial species in natural ecosystems (Garud *et al*. 2019; Ramiro *et al*. 2020). In particular, Garud et al. (2019) reported that the purifying selection for protein sequences within species is much weaker than that between species, suggesting a cautionary note for the applicability of the E–R anticorrelation in relatively recent evolution among bacteria. In addition, recent studies have also pointed out the inconsistency between diverse empirical data across multiple organisms and the predictions from the frequently suggested possible mechanisms explaining the E–R anticorrelation (Plata *et al*. 2010; Plata and Vitkup 2018; Razban 2019; Usmanova *et al*. 2021). For instance, recent genome-scale data empirically measuring protein stability, protein aggregation, and protein stickiness do not support the considerable extent of selection against protein misfolding or protein misinteraction for highly expressed proteins in *Escherichia coli* (Usmanova *et al*. 2021). In turn, these conflicts raise questions about the generality of the E–R anticorrelation and the relevance of the plausible mechanisms governing it, which motivated us to test the applicability of the E–R anticorrelation on bacterial microevolution and the relevance of the possible underlying mechanisms.

To this end, we analyzed the genome sequences of 99 strains of *E. coli*, whose mutations accumulated through microevolution in nature. We also explored the E–R anticorrelation of *de novo* evolution via an evolution experiment using *E. coli*. We found significant E–R anticorrelation in both past and *de novo* evolution in *E. coli*. We also found that purifying selection acting on highly expressed genes contributed to the ubiquity of the E–R anticorrelation. This study confirmed that purifying selection acting on highly expressed genes is not an evolutionary legacy but rather an active component, implying that expression level has a ubiquitous impact on the speed of evolutionary molecular diversification in bacteria. The detected selection included codon-level purifying selection, which supports the relevance of the underlying mechanisms proposed previously. Nevertheless, their effects on recent evolution may be smaller than expected. Our study emphasizes the importance of the expression level in understanding how genetic divergence emerges within a bacterial species and also provides new insight into the controversy of the dominant mechanisms underlying the E–R anticorrelation.

## Results

### The inter- and intraspecific E–R anticorrelation in past evolution

The rate of interspecific evolution among protein sequences can be explained by the interrelationship between the number of nonsynonymous nucleotide changes per nonsynonymous site (dN) and the number of synonymous nucleotide changes per synonymous site (dS) in the orthologous genes between closely related species (Figure 1A). We refer to interspecific dN and dS as dN_btw_ and dS_btw_, respectively. Previous studies have shown that both dN_btw_ and dS_btw_ are negatively correlated with expression levels in *E. coli* (Figure 1B, C) and in other organisms (Drummond and Wilke 2008). The underlying mechanisms of these relationships are explained by purifying selection at the codon level (Drummond and Wilke 2008; Yang *et al*. 2010; Park *et al*. 2013). In particular, the protein misfolding avoidance hypothesis (Yang *et al*. 2010) explains that optimal codons are favored in highly expressed proteins to avoid toxic misfolding and that dN_btw_ and dS_btw_ are common rather than independent targets of codon-level purifying selection to combat misfolding. Consistent with this hypothesis, we found a negative correlation between dN_btw_/dS_btw_ and the expression level. The correlation was somewhat weaker than the E–R anticorrelation in dN_btw_, most likely due to the fact that the common purifying selection acting on dN_btw_ and dS_btw_ was cancelled out (Figure 1D). Nevertheless, the negative correlation between dN_btw_/dS_btw_ and the expression level remains substantial, suggesting that another mechanism contributes to purifying selection which acts on highly expressed genes.

**Figure 1.**
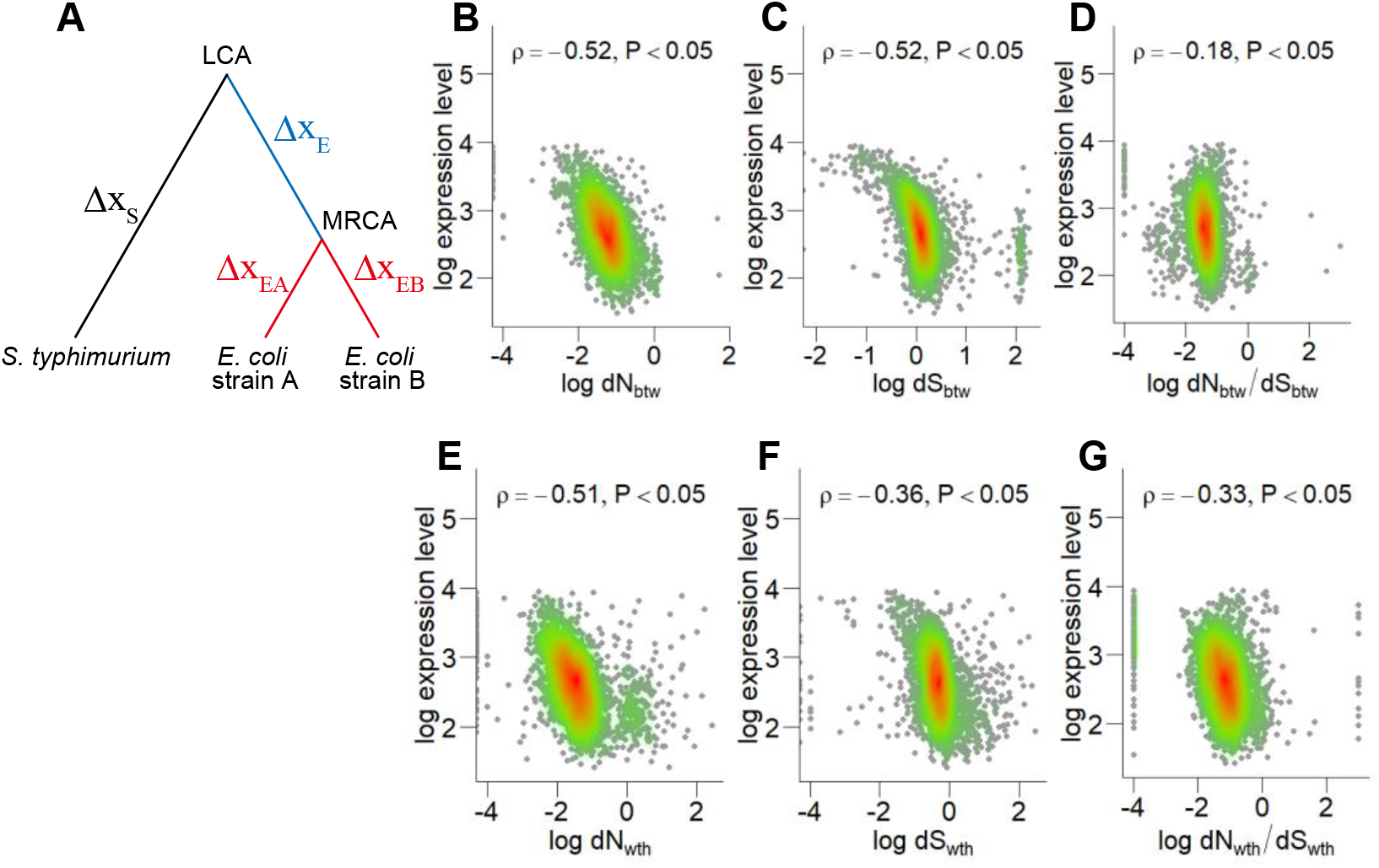
The negative correlation between mRNA expression level and the rates of DNA sequence of orthologs in the course of past evolution. (A) A schematic phylogeny of *E. coli* and *Salmonella typhimurium*. Genetic changes between nodes are indicated as Δx_S_ for *S. typhimurium* from the last common ancestor of *E. coli* and *S. typhimurium* (LCA), Δx_E_ for the most recent common ancestor of *E. coli* (MRCA) from the LCA, Δx_EA_ and Δx_EB_ for *E. coli* strain A and B from the MRCA, respectively. Genetic changes between species, N_btw_ and S_btw_ included, represent the difference between Δx_S_ and the sum of Δx_E_ and Δx_EA_ or the sum of Δx_E_ and Δx_EA_. Genetic changes within *E. coli* species, N_wth_ and S_wth_ included, represent differences between Δx_EA_ and Δx_EB_. (B–D) The negative correlation of the rate of interspecific evolution of DNA sequences (*E. coli* and *S. typhimurium*). (E–G) The negative correlation of the rate of intraspecific evolution of DNA sequences (*E. coli*). The evolutionary rate of the DNA sequence is characterized by dN (B, E) and dS (C, F), respectively. (D, G) The dN/dS ratio of interspecific (D) and intraspecific evolution (G). The expression level was calculated from *E. coli* transcriptome data. Each dot corresponds to a single gene. The red-green gradient represents the 2D density (high to low). Spearman’s rank correlation coefficients and p-values are shown.

To test whether within-species molecular evolution also follows the E–R anticorrelation, we quantified intraspecific dN and dS, referred to as dN_wth_ and dS_wth_, among 99 strains of *E. coli*. We found that both dN_wth_ and dS_wth_ were negatively correlated with gene expression relative to that of interspecific evolution (Figure 1E, F). In addition, the correlation coefficient for dN_wth_ was slightly larger than that for dS_wth_, which was similar to the genetic signatures of interspecific evolution in other organisms, such as yeast or flies. This difference between dN_wth_ and dS_wth_ also suggests that the E–R anticorrelation in dN_wth_ reflects different purifying selections from those acting on dS_wth_, as in the case of the E–R anticorrelation in dN_btw_. To confirm this hypothesis, we explored the relationship between dN_wth_/dS_wth_ and expression levels. As with the case of interspecific evolution, dN_wth_/dS_wth_ showed a substantial negative correlation with expression level, although the correlation was weaker than the E–R anticorrelation in dN_wth_. Therefore, the purifying selection on dS_wth_ seems to be insufficient to explain the E–R anticorrelation in intraspecific evolution. These results suggest that E–R anticorrelation itself might be causal to a general pattern of molecular evolution in the past, but the underlying mechanisms of purifying selection remain an open question, as stated recently in the literature (Plata and Vitkup 2018).

### E–R anticorrelation in *de novo* evolution

To determine whether the E–R anticorrelation is an evolutionary legacy or is currently applicable, we explored the relationship between protein evolutionary speed and gene expression levels during *de novo* evolution. Using a previously developed UV-irradiating cell culture device (Shibai *et al*. 2019), we conducted an evolution experiment to rapidly accumulate mutations (Figure 2A). *Escherichia coli* cells were incubated in this device and transferred to a fresh medium every four days. During incubation, the device automatically measured the optical density (OD) of the culture and irradiated UV for each unit increment of OD, where UV was utilized as a mutagen and germicidal lamp (Figure 2B). This feedback control of UV irradiation prevented the depression of mutation rates caused by the acquisition of UV resistance in the cells. We established six independent lineages from an ancestral colony and repeated the cycle of incubation and transfer for two years, corresponding to tens of thousands of generations (Figure 2C). As a result, we obtained thousands of base-pair substitutions (BPSs) of the coding region fixed in each cell population (Figure 2D). The occurrence of the same mutations over multiple lineages was exceedingly rare, ensuring that most of the accumulated BPSs contributed to the evolutionary diversification of the DNA sequence. To understand the overall evolutionary processes of diversification, we calculated whole-genome dN/dS values (Figure 2E) by considering a mutational spectrum (Figure 2F). The dN/dS of most lineages was roughly 0.9, indicating that most BPSs were fixed in the populations through neutral processes rather than by adaptive processes. Moreover, considering the large population size and high mutation rate in the culture device, many of these non-synonymous BPSs were likely to be fixed in the population by hitchhiking rather than genetic drift.

**Figure 2.**
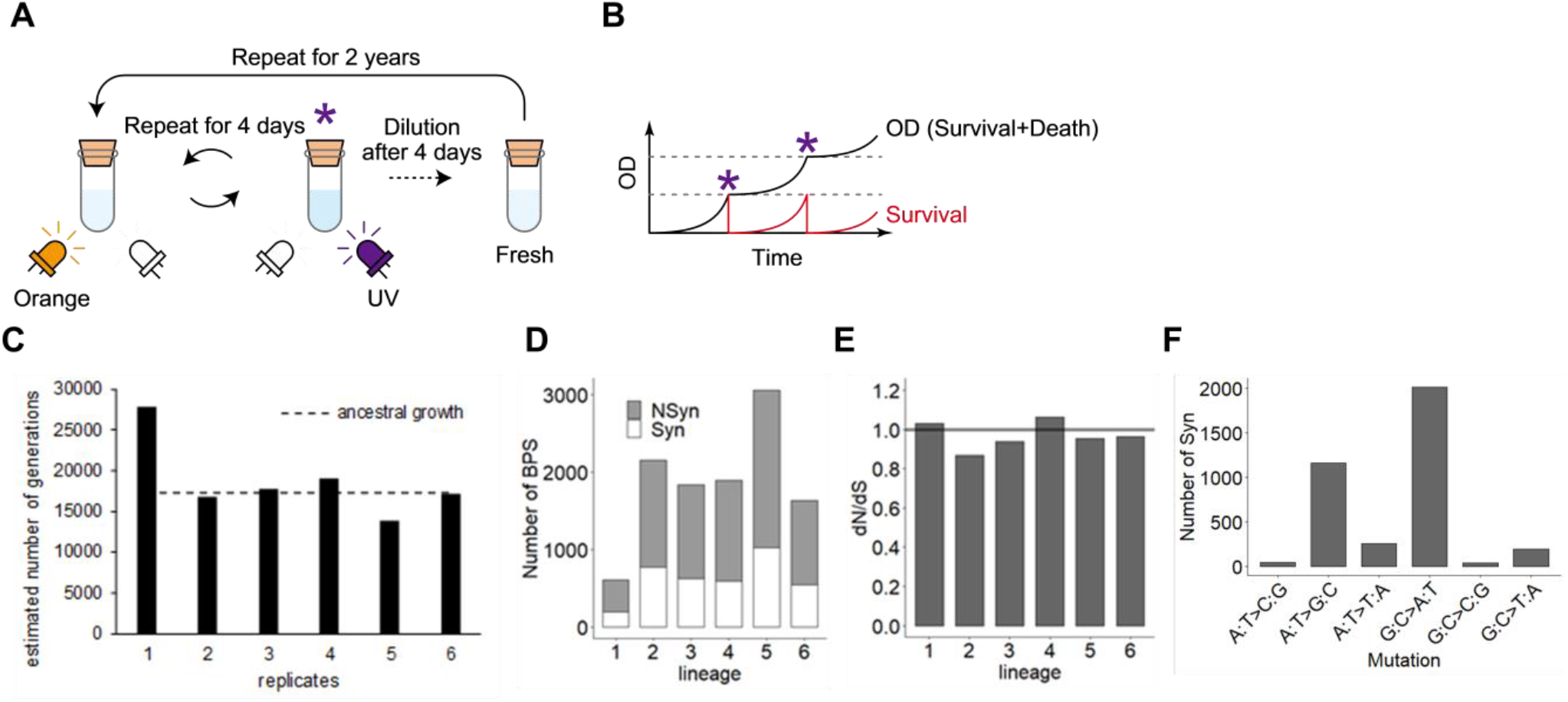
Evolution experiment for accumulating massive mutations. (A) Procedure of an evolution experiment with the UV irradiating cell culture device. The device consists of a quartz glass test tube with a resin housing that measures the cell density (OD) by an orange LED and irradiates UV light by a UV-C LED. Mutagenesis by UV irradiation (denoted as purple asterisks) was performed when OD exceeded a defined increment so that the survival fraction could be maintained within a constant range (B). After four days of repeats, an aliquot of cell culture was diluted with fresh media 100 times and transferred into a new test tube. These procedures were repeated for six independent replicates for two years. (C) The estimated number of generations after 688 days of the evolution experiments. The black bars correspond to the values calculated with the doubling time of evolved cells for each of the six replicates. The dashed line indicates the value calculated with the ancestral doubling time. (D) The number of accumulated BPSs during the evolution experiment. The gray and white fractions of a bar represent nonsynonymous and synonymous substitutions, respectively. (E) The genome-wide dN/dS values were close to 1.0 for all the six replicates, implying that the majority of the accumulated mutations had neutral effects on their fixation within the populations. (F) Mutation spectrum of synonymous substitutions. The synonymous substitutions of all lineages are summed for each substitution type.

To explore the expression levels of the mutated genes, we obtained transcriptome data of the ancestral and evolved samples by microarray and quantified the geometric mean of six independent lineages. We found that the expression profiles of the evolved strains were similar to that of the ancestral strain (ρ = 0.89–0.94, Figure S1). Using transcriptome data, we explored the relationship between the protein evolutionary rate and gene expression levels during *de novo* evolution. For each gene, we quantified dN and dS in *de novo* evolution, referred to as dN_novo_ and dS_novo_, by using the sum of the number of nonsynonymous and synonymous BPSs among six independent lineages. We found a significant E–R anticorrelation even in *de novo* evolution, regardless of ancestral (ρ = −0.17, p < 0.05) or evolved expression levels (ρ = −0.19, p < 0.05, Figure 3A). We also confirmed that this negative correlation remained after controlling for gene dispensability (ρ = − 0.18, p < 0.05, partial correlation test for maximal growth rate of deletion mutants). Notably, the mutation data of approximately half the number of total mutations (i.e., the data at one-year of evolution) exhibited a similar negative correlation (ρ = −0.16, p < 0.05). Thus, we confirmed that the observed E–R anticorrelation was relatively weak but insensitive to the progress of our evolution experiment or to changes in transcription profiles, at least during our evolution experiment. Contrary to the evolution between species, the negative correlation between dS_novo_ and expression levels was found to be much weaker than that of dN_novo_ (Figure 3B). We also confirmed a negative correlation between dN_novo_/dS_novo_ expression levels (Figure 3C), as well as the E–R anticorrelation in dN_novo_. Thus, our *de novo* evolution experiments revealed ongoing purifying selection acting on highly expressed genes.

**Figure 3.**
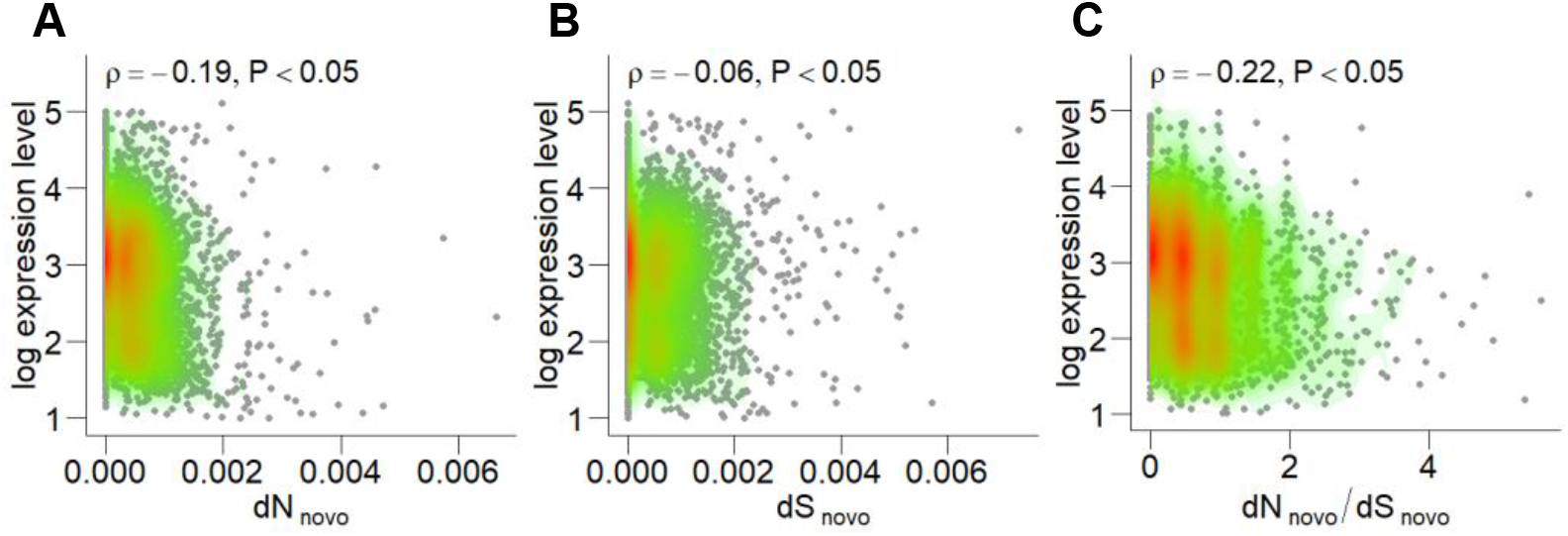
There was a negative correlation between the protein sequence evolution during the evolution experiment and the gene expression level. (A) dN_novo_ showed a negative correlation with the gene expression level. (B) On the other hand, dS_novo_ showed only a slight correlation with the expression level. (C) A negative correlation was also observed for dN_novo_/dS_novo_, where dN_novo_ was normalized by dS_novo_ by cancelling the common selection.

### Purifying selection on codon usage in *de novo* evolution was less sensitive to expression level

The expression level dependency of dS reflects the purifying selection of codon usage of highly expressed proteins, which is a frequently suggested explanation for the E–R anticorrelation in dN (Drummond and Wilke 2008). Highly expressed proteins use optimal codons that enable fast and accurate translation (Akashi 2001, 2003) and protein stability (Yang *et al*. 2010). The use of other unfavorable codons has detrimental effects on cellular growth and is thought to be evolutionarily constrained (Zhang and Yang 2015). However, the small anticorrelation between dS_novo_ and expression levels obscures the expected expression dependency of the purifying selection on codon usage in *de novo* evolution. To clarify this, we explored the relationship between the degree of codon optimization of each protein and the evolutionary speed of synonymous BPSs. Since this relationship is expected to be weak, it is important to evaluate the evolutionary speed of a small number of synonymous BPSs. To this end, we used a normalized version of the G score, hereinafter referred to as the G score, as an alternative to dN_novo_ and dS_novo_, as detailed in the **Materials and Methods**. The G score is useful for screening genes with a small number of substitutions relative to neutral expectations. First, we reconfirmed the E–R anticorrelation between expression level and G score in nonsynonymous substitutions (G_N_, ρ = −0.15, p < 0.05) and that there was no correlation in synonymous substitutions (G_S_), which was consistent with the relationship between expression level and dN_novo_ or dS_novo_. Next, we employed the codon adaptation index (CAI) as a standard measure of the degree of codon optimization and explored the relationship between CAI and G scores. As a result, a negative correlation was found between the CAI and G score for nonsynonymous BPSs (Figure 4A) and synonymous BPSs (Figure 4B), although the correlation coefficient for synonymous BPSs was not strong. To confirm the looseness of the purifying selection on codon-optimized proteins in *de novo* evolution, we classified 10% of mutated proteins with the lowest CAI as unoptimized, 10% of mutated proteins with the highest CAI as optimized, and the remaining mutated proteins as having moderate optimality in terms of codon usage for nonsynonymous and synonymous BPSs. As expected, unoptimized proteins showed higher G_S_ than the optimized and moderately optimized proteins (Figure 4D). In contrast, there was no significant difference between optimized and moderately optimized proteins, indicating that the purifying selection on codon usage only weakly depends on expression levels in *de novo* evolution. This tendency remained even if the classification criteria for CAI changed from 10% to 5%. To confirm the looseness of the purifying selection on codon usage more directly, we focused on individual synonymous BPSs and explored codon bias. To this end, we calculated the C score for synonymous BPSs, whereby the C score represents the difference in preference of the mutant synonymous codon from neutral expectation, as detailed in the **Materials and Methods**. In short, the C score takes positive values if the mutant synonymous codons are used more frequently in highly expressed proteins than in neutral expectations, while it takes negative values if the mutant synonymous codons are used less frequently in highly expressed proteins than in neutral expectations. Contrary to the statistics, such as G scores or CAI, characterizing each gene, C scores are assigned to each synonymous BPS, not to each gene. In other words, each gene had as many C scores as the number of synonymous BPSs in each gene. We found that unoptimized proteins allowed for more mutant synonymous codons, which are infrequently used in highly expressed proteins than moderately optimized codons (Figure 4E). In contrast to the other categories, the mutant synonymous codons of the optimized proteins were not able to obtain high C scores because the wild-type codons of the optimized proteins are likely to be the most frequent among the highly expressed proteins. Therefore, it is rational that there was no statistical significance between optimized and unoptimized proteins, even though the C score of the former was relatively larger than that of the latter. Altogether, these results support that the detected purifying selection on codon usage is active but less sensitive to expression levels.

**Figure 4.**
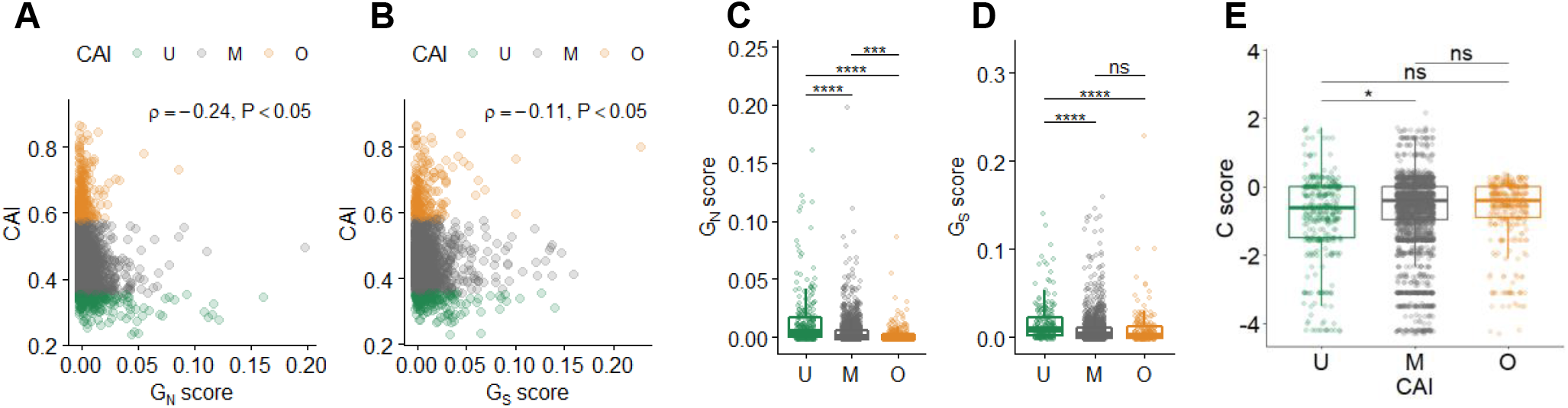
Relation between G scores and Codon Adaptation Index. The Codon Adaptation Index (CAI) was negatively correlated with G scores for nonsynonymous (G_N_, A) and synonymous BPSs (G_S_, B). Spearman’s rank correlation and p-values are indicated in each panel. Color represents codon optimality (U, unoptimized; M, moderate; O, optimized proteins). Comparison between codon optimality and G scores (G_N_, C; G_S_, D). Enlarged panels are shown at the bottom. (E) Comparison between codon optimality and C score. Adjusted p-values for Wilcoxon test are indicated as ns > 0.05, * < 0.05, ***< 0.001, and **** < 0.0001.

### Purifying selection of synonymous substitution on molecular function

The difference between dN_novo_ (or G_N_) and dS_novo_ (or G_S_) in correlation with expression levels suggests that the protein features on which purifying selection acts in *de novo* evolution of synonymous BPSs might be somewhat different from that of nonsynonymous BPSs. To confirm this possibility, we conducted a GO enrichment analysis for the proteins ranked in the top or bottom 10% of G scores for synonymous and nonsynonymous BPSs (Figure 5). We found 70 GO terms enriched in the bottom 10% of G_S_; in contrast, no GO terms were enriched in the bottom 10% of G_N_ (Figure 5A). Interestingly, all of the enriched terms were classified in the molecular function category, suggesting that some enzymatic features were related to the target of purifying selection for synonymous BPSs rather than any metabolic pathways. For instance, the enriched GO terms contained ATPase activity (GO:0016887), which is required for various biochemical reactions (Figure 5D), regardless of metabolic pathways. Contrary to the bottom 10% of G_S_, the top 10% of G_S_ showed no enrichment in the molecular function category; however, 17 GO terms were enriched in the biological process category, such as the lipopolysaccharide biosynthetic process (GO:0009103). Many of these were common among the GO terms enriched in the top 10% of G_N_ (Figure 5B, C), suggesting that some proteins related to these processes were likely to be inactivated and were not targeted by purifying selection for both synonymous and nonsynonymous BPSs. These results support the hypothesis that the purifying selection acting on synonymous BPSs is not a single dominant mechanism of purifying selection on nonsynonymous BPSs, at least in *de novo* evolution.

**Figure 5.**
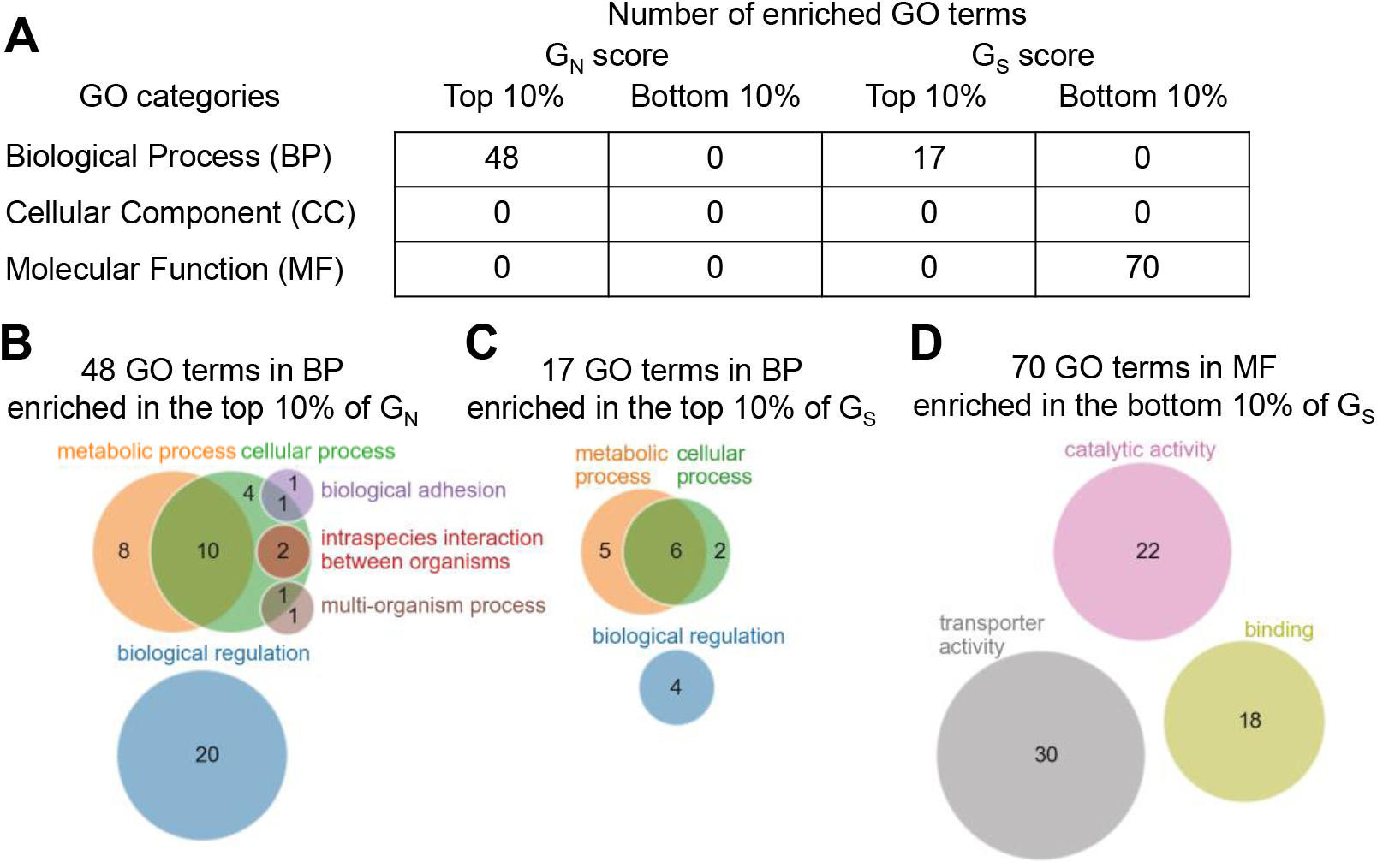
Comparison between G scores with biological features. (A) Enrichment analysis for the top and bottom 10% of G_N_ and G_S_. The number of GOs enriched significantly was shown in each class. (B–D) Venn diagram of the ancestral GOs at the second level (circles) of the GO tree for each of the enriched GOs (B for top 10% of G_N_, C for top 10% of G_S_ and D for bottom 10% of G_S_). The number of enriched GOs in each parental GO is indicated in each circle. BP, Biological Process; MF, Molecular Function.

## Discussion

The present study explored the impact of expression levels on the molecular evolution of bacteria. By employing comparative genomics and a laboratory-based evolution experiment, we elucidated the ubiquity of the impact of expression level on the evolutionary speed of sequence diversification. We found that the E–R anticorrelation governs not only sequence diversification between species but also within species. This finding of the ubiquity of the E–R anticorrelation is consistent with the recent analysis of genomic mutations accumulated in *E. coli* over long-term evolution experiments (Maddamsetti 2021). However, there are several disparities between the latter and the present study. First, the correlation coefficients between the expression level and the rate of nonsynonymous mutations in the long-term evolution experiments were almost negative, but their magnitudes were much smaller (ρ = −0.0486––0.0991) than those for *de novo* evolution in our study (ρ = −0.19, Figure 3A). Second, the correlation coefficients between the expression level and the rate of synonymous mutations in the long-term evolution experiments were positive (0.0458–0.094), contrary to the negative values in our *de novo* evolution experiment (ρ = −0.06, Figure 3B) and natural microevolution (ρ = −0.36, Figure 1F). We speculate that these differences arose not only from the difference in conditions between the two evolution experiments, but also from the difference in the analytical method used to calculate the evolutionary speeds of DNA sequences. Contrary to our study, for example, the previous study included mutations unfixed in the populations to calculate the evolutionary speeds. Accounting for unfixed mutations tends to obscure the signatures of natural selection and is likely to underestimate purifying selection. In addition, the previous study did not use dN or dS but rather employed the number of nonsynonymous or synonymous mutations per length as a measure of the rate of evolution. Accordingly, neither biased mutational spectrum nor the differences in probability of synonymous/nonsynonymous sites among genes were considered properly, which could interfere with the calculation of the evolutionary speeds for each gene. On the other hand, our method carefully treats these key factors when measuring the evolutionary speeds of DNA sequences, as detailed in the **Materials and Methods**. Thus, our data support the reliability of the E–R anticorrelations. We also found that the purifying selection acting on highly expressed genes is not a legacy but actively constrains the sequence diversification of these genes, even along a relatively short evolutionary timescale. The detected selection included purifying selection at the codon level, supporting the relevance of the possible underlying mechanisms such as selection against protein misfolding or protein misinteraction, since these frequently suggested mechanisms assert codon-level purifying selection acting on highly expressed proteins (Yang *et al*. 2010, 2012). Nevertheless, our data also suggest that the impacts of these frequently suggested possible mechanisms on recent evolution might be weaker than previously expected. These findings are consistent with recent studies indicating that empirical data measuring protein stability, protein aggregation, and protein stickiness do not support the considerable impact of these frequently suggested mechanisms on the E–R anticorrelation for macroevolution (Plata *et al*. 2010; Plata and Vitkup 2018; Razban 2019; Usmanova *et al*. 2021). Therefore, the unexpected weak impacts of the frequently suggested mechanisms might be common between macro-and microevolution. In conclusion, this study suggests the importance of the expression level when attempting to understand how genetic divergence emerges within a bacterial species, and also provides a new insight into the controversy of the dominant mechanisms underlying the E–R anticorrelation (Zhang and Yang 2015).

In this study, the E–R anticorrelation was observed in both past and *de novo* microevolution. However, the negative correlation of the former is stronger than that of the latter. What does this difference mean? We speculated that the magnitude of purifying selection against protein sequences could explain this difference, since the E–R anticorrelation mainly reflects the purifying selection. We found this to be true. In our experiment, the average dN/dS of past evolution was smaller than that of *de novo* evolution. That is, purifying selection against protein sequences in past evolution is stronger than that of *de novo* evolution. Why is the purifying selection in *de novo* evolution relatively small, even in the presence of selection for growth/survival in our evolution experiment? There are at least two plausible explanations for this finding. The first possible and trivial explanation is that natural environments are more severe than those experienced in test tubes. Under our laboratory conditions, the nutrients required for growth were supplied constantly and at sufficient levels. In addition, the stress factor was limited to that from the UV alone. On the other hand, the quality and quantity of both nutrients and stressors must be different from the laboratory conditions and must change unpredictably. These severe conditions enable us to speculate that the essentiality of each gene is strong even for nonessential genes, which are characterized in relatively milder laboratory conditions. In other words, the detrimental effects of a given mutation are strong under natural conditions. Therefore, it is not difficult to imagine that a strong purifying selection governs evolution in nature. The second explanation is plausible if we consider a high mutation rate in our evolution experiment. The rate of mutation in our experimental setup was hundreds of times higher than the spontaneous mutation rate that would be experienced in nature. Therefore, neutral-to-deleterious mutations are relatively frequent. The population bottleneck in our experiment was large enough to fix these frequent deleterious mutations in a population by hitchhiking driver beneficial mutations. Therefore, the deleterious effects of a given passenger mutation are alleviated by the beneficial effects of driver mutations. As a result, purifying selection cannot purge such alleviated detrimental mutations, which yields nearly neutral values for dN/dS. These mechanisms are non-mutually exclusive. Interestingly, a high mutation rate and neutrality driven by hitchhiking are not only applicable to our artificial condition, but are also seen in more natural situations (Ramiro *et al*. 2020). Therefore, the relaxation of purifying selection due to high mutation rates may partially contribute to past divergent evolution within species.

Why is the E–R anticorrelation regarded as being general? The mechanical origins of the E–R anticorrelation have been extensively proposed, such as the protein misfolding avoidance hypothesis or the misinteraction avoidance hypothesis. However, most of the proposed mechanisms cannot fully explain the generality of the E–R anticorrelation. Previous studies have focused on identifying the type of fundamental biological processes for a mutated gene that has deleterious effects on any organism. In contrast, our results suggest the importance of robustness or conservativeness of the entire transcriptional expression pattern during evolution to explain the generality of the E–R anticorrelation. If expression levels evolve without any constraints or are highly dynamic, the E–R anticorrelation would lose its generality. The expression level of a gene is expected to change dynamically during evolution, for example, by the mutation of a corresponding transcription factor or intergenic region. In fact, an enrichment analysis detected those nonsynonymous mutations significantly accumulated transcription factors in our evolution experiment. Interestingly, however, the entire transcription level exhibited only slight changes from the ancestor even after the accumulation of thousands of mutations. As a result, an equivalent level of the E–R anticorrelation was observed in both the ancestral transcriptional data and in the evolved transcriptional data (rho = −0.21~–0.23). Such conservativeness among expression levels was also detected in other evolutionary experiments equipped with growth selection. For example, Ho and Zhang (2018) revealed that genetic changes more frequently reverse rather than reinforce transcriptional plastic changes in adaptation to a new environment, generally because an original transcriptional state is favored during growth selection. Transcriptome level conservation has also been observed in bacterial evolution in nature (Zarrineh *et al*. 2014; Payne and Wagner 2015; Junier and Rivoire 2016). Likewise, any compensatory mutations might restore expression levels that were altered by other harmful mutations to their original levels in our evolution experiment. Therefore, some mutations among transcriptional factors may play a role in compensatory mutations to retain their expression levels. In addition to the genetic mechanism, there are cases in which an alternative mechanism without any mutations underlies conservativeness at the expression level. For instance, Briat et al. (Briat *et al*. 2016) proposed a network motif conferring homeostasis or the perfect adaptation of expression levels to intrinsic and extrinsic disturbances. Such mechanisms are also applicable to mutational disturbances in the expression levels. In addition, it has been pointed out that ORFs can somehow determine their own expression levels (Isalan *et al*. 2008). To understand the generality of the E–R anticorrelation, the present study sheds light on the importance of understanding the quantitative relationship between protein sequence evolution and expression evolution.

## Materials and Methods

### Database analysis of mRNA expression levels

A total of 218 microarray datasets of *E. coli* K-12 substrain MG1655 with the GPL3154 platform were used in this study (Table S1). They were included in 27 experiments and downloaded from the Gene Expression Omnibus (Barrett *et al*. 2013). After quantile normalization (Bolstad *et al*. 2003), the average and variance of the expression levels were calculated for each gene.

### Interspecific analysis of protein evolution

The protein evolutionary rates of *E. coli* were obtained from the literature, which compared the genomes of *Escherichia coli* K-12 MG1655 and *Salmonella typhimurium* LT2 (“Supplementary information S2” in Zhang and Yang 2015). The dN and dS values were calculated from the genomic sequences of *E. coli* str. K-12 substr. MG1655, and *Salmonella enterica* subsp. *enterica* serovar Typhimurium str. LT2 (accession no. NC_000913.3 and NC_003197.2). A total of 3145 paired sets of orthologous genes were detected by the bidirectional best hits (Overbeek *et al*. 1999) method, comparing all combinations of two coding features from the genomes. For each orthologous gene set, Clustal Omega (McWilliam *et al*. 2013) was used to generate an alignment, and PAML was used to calculate the dN and dS values from the alignment (Yang 1997).

### Intraspecific dN/dS analysis

The coding DNA sequences for 99 *E. coli* genomes were downloaded from Ensembl Genomes (Zerbino *et al*. 2018) in the multi-fasta format (Table S2). Each coding feature of the genomes was annotated by the bidirectional best hits (Overbeek *et al*. 1999) method compared with *the E. coli* K-12 substrain MG1655, generating groups of orthologous genes. Clustal Omega and Clustal W2 (McWilliam *et al*. 2013) aligned the sequences and generated phylogenetic trees for each orthologous group. PAML (Yang 1997) calculated the dN and dS values for each tree.

### Strain and culture conditions

We used the *E. coli* K12 substrain MDS42 (Pósfai *et al*. 2006) as the ancestor of the evolution experiment. We used a chemically defined medium, mM63, which comprised 62 mM K_2_HPO_4_, 39 mM KH_2_PO_4_, 15 mM (NH_4_)_2_SO_4_, 2 μM FeSO_4_·7H_2_O, 15 μM thiamine hydrochloride, 203 μM MgSO_4_·7 H_2_O, and 22 mM glucose (Kashiwagi *et al*. 2009). The cells were inoculated into 8 mL of the mM63 medium and incubated with shaking at 37 °C.

### Evolution experiment

The evolution experiment procedure consisted of a 4-day cycle of a serial transfer cycle. We used an automated UV-irradiating cell culture system that was previously reported (Shibai *et al*. 2019). First, the OD value of the cell culture was measured automatically. When the OD value exceeded the stipulated threshold (OD_THR_), the cells were exposed to a dose of UV light which killed the cells, resulting in a survival rate of the ancestral cell of 10^−2^ to 10^−3^. Then, the threshold, OD_THR_, was renewed as OD_THR_+OD_STEP_, so that the next UV irradiation was conducted when the living cell population recovered to the amount corresponding to OD_STEP_. The OD_STEP_ and initial OD_THR_ values were set at OD_600_ = 0.0015. The cells were glycerol-stocked at the end of each round.

### Whole-genome resequencing

Cells were grown in a mM63 medium at 37 °C with shaking at 200 rpm overnight for 2 days, which were then pelleted by centrifugation. Genomic DNA was extracted from the cells using a Wizard Genomic DNA Purification Kit (Promega). DNA libraries were prepared using a Nextera XT kit (Illumina) for paired-end sequencing (2× 300 bp), according to the manufacturer’s instructions. Illumina MiSeq sequenced the DNA libraries using the MiSeq Reagent Kit v3 for 600 cycles. Mutation detection was performed by mapping the resulting read data to the reference genome sequence (accession no. AP012306.1) using the Burrows-Wheeler Aligner software (Li and Durbin 2009) and SAMtools (Li *et al*. 2009). For quality control, the called mutations were filtered using the Phred quality score (Ewing and Green 1998; Cock *et al*. 2009) with a cut-off value of > 100. In addition, base-pair substitutions (BPSs) with a frequency of “mutant” reads < 90% were removed. The resulting mutations were annotated using an in-house program written in C++.

### Calculation of dN and dS in *de novo* evolution

Genome-wide dN/dS values were calculated from the numbers of both synonymous and nonsynonymous BPSs using a previously reported method (Shibai *et al*. 2017). dN and dS values in *de novo* evolution for each gene, referred to as dN_novo_ and dS_novo_, were calculated similarly, assuming each gene sequence as a full-length sequence.

### Calculation of G scores

The G score was defined as the actual number of mutations (M) multiplied by the logarithm of the ratio of the actual number of mutations to the expected number of mutations (log(M/E)) (Tenaillon *et al*. 2016). Therefore, the G score was supposed to show positive values with mutationally accelerated genes, negative values with suppressed genes, and zero values with non-biased genes. In this study, we normalized the G score by the number of mutational sites in each gene for more precise bias analyses. Specifically, the G score of each gene for synonymous (subscripted with *S*) and nonsynonymous (subscripted with *N*) substitutions were calculated according to the following formulas:

Normalized G score of synonymous and nonsynonymous substitutions of gene *i*:

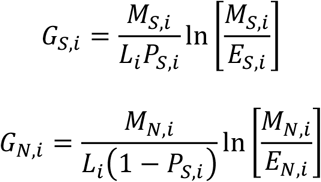

Expected number of synonymous and nonsynonymous substitutions of gene *i*:

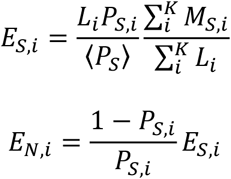

*M*_*S,i*_: observed number of synonymous substitutions in gene *i*

*M*_*N,i*_: observed number of nonsynonymous substitutions in gene *i*

*K*: number of genes in the genome

*L*_*i*_: length of the coding DNA sequence of gene *i*

*P*_*S,i*_: the probability that the substitution is synonymous substitution when a substitution occurs in gene *i* as detailed below. ⟨*P*_*S*_⟩ represents the mean of *P*_*S,i*_ for all the genes.

The probability that the substitution occurred on a given codon when a substitution occurred in gene *i* was calculated using the following equation:

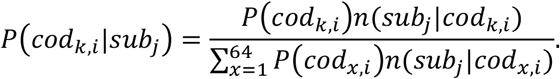

Here, each substitution of all six possible substitutions is denoted by *sub*_*j*_, where *j* takes 1–6, using the following array:

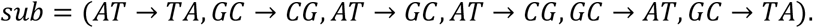

In addition, each codon of all 64 possible codons in a given gene *i* is denoted by *cod*_*k,i*_, where *k* takes 1–64, using the following array:

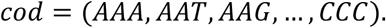

The codon usage of codon *k* in gene *i* is then represented by *P*(*cod*_*k,i*_), which was calculated from the genome sequence of the ancestral strain. In addition, the number of possible mutant triplets when the *j* th substitution occurs in a given *cod*_*k*_ in gene *i* is denoted by *n*(*sub*_*j*_|*cod*_*k,i*_). Therefore, the probability of synonymous change for a given codon in gene *i* with a given *j* th substitution is given by the following equation:

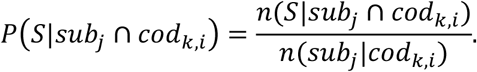

Here, the number of synonymous triplets when a *sub*_*j*_ occurs in a given *cod*_*k,i*_ is denoted by *n*(*S*|*sub*_*j*_ ∩ *cod*_*k,i*_). Using the mutational spectrum for synonymous substitutions, *P*(*sub*_*j*_), these two probabilities give *P*_*S,i*_ using the following equation:

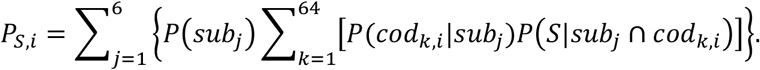

### Calculation of the Codon Adaptation Index

The codon adaptation index (CAI) indicates the abundance of optimal codons in a gene sequence, where an optimal codon is defined as the most frequent codon in each of the synonymous codon groups used in the most abundant proteins (Sharp and Li 1987). The CAI of a given gene with an amino acid length *La* was calculated as follows:

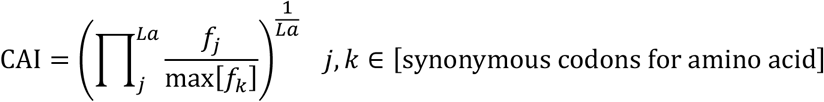

where *f*_*j*_ is the frequency of the codon coding for *j*th amino acid of the given gene and max [*f*_*k*_] represents the frequency of the most frequent synonymous codon *f*_*k*_ for that amino acid. We calculated the frequency of each codon by considering the 40 most abundant genes based on the transcriptome of the ancestral strain.

### Calculation of C score

The C score is an indicator of bias in codon weight change caused by a synonymous substitution. Note that the C score was calculated for each mutation, not for each gene, as in the other indicators used in this study. The C score in which an ancestral codon (*a*) changes to a mutated codon (*m*), referred to as *C*_*a*→*m*_, is defined as follows:

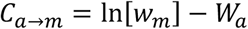

where *w*_*m*_ is the codon weight of codon *m* calculated by the following formula:

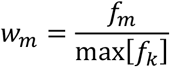

where *f*_*m*_ is the frequency of codon *m* of the focal amino acid and max [*f*_*k*_] represents the frequency of the most frequent synonymous codon *f*_*k*_ for that amino acid. In addition, *W*_*a*_ is the average of the logarithms of the codon weights with a single synonymous substitution of codon *a*, and corresponds to the expected value of the mutated codon weights as follows:

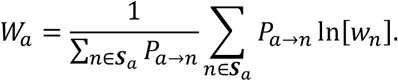

***S***_*a*_: set of all possible synonymous codons from a given ancestral codon *a* by a single BPS. *m ∈* ***S***_*a*_.

*P*_*a*→*n*_: frequency of a BPS that enables synonymous mutation from codon *a* to codon *n*, which was calculated by the mutational spectrum of synonymous substitutions.

### Gene ontology analysis

Gene ontology (GO) enrichment analysis was performed using GOstats (v.2.48.0, R Bioconductor) (Falcon and Gentleman 2007). We used all three categories: biological process (BP), molecular functions (MF), and cellular components (CC). The resulting GO terms were filtered with cutoffs of 0.01 and 0.05 for their respective p-value and q-value (Storey *et al*. 2021). Genes within the top and bottom 10% of the normalized G score were analyzed as gene sets. For visualization, the detected GO terms were converted to their ancestral GO terms in the second level of the gene ontology tree, that is, the layers directly under BP, MF, or CC.

### mRNA expression profiling of genes using microarray technology

The cells were cultured for 16–19 h and then sampled at the time of the logarithmic growth phase (OD_600_ values were 0.072–0.135). Aliquots of the cells were immediately added to the same volume of ice-cold ethanol containing 10% (w/v) phenol. RNA extraction was performed using a RNeasy mini kit with on-column DNase digestion (Qiagen), following the manufacturer’s protocol. The purified RNA was quality-controlled using an Agilent 2100 Bioanalyzer and an RNA 6000 Nano kit (Agilent Technologies). A microarray experiment was performed using an Agilent 8× 60 K array, which was designed for the *E. coli* W3110 strain so that 12 probes were contained for each gene. Purified total RNA (100 ng) was labelled with Cyanine3 (Cy3) using a Low Input Quick Amp WT labelling kit (One-color; Agilent Technologies). The Cy3-labelled cRNA was checked for its amount (> 5 μg) and specific activity (> 25 pmol/μg) using NanoDrop ND-2000. Then, the cRNA of 600 ng was fragmented and hybridized to a microarray for 17 h at 65 °C, rotating at 10 rpm in a hybridization oven (Agilent Technologies). The microarray was then washed and scanned according to the manufacturer’s instructions. Microarray image analysis was performed using Feature Extraction version 10.7.3.1 (Agilent Technologies). The resulting gene expression levels were normalized using quantile normalization.

## Data Availability

The raw sequence data of genome sequence analyses the ancestral and evolved samples in this article are available in NCBI’s Sequence Read Archive (SRA) under the accession numbers SRR16961197 to SRR16961208. The microarray data of the ancestral and evolved samples in this article are available in NCBI’s Gene Expression Omnibus (GEO) and are accessible through GEO Series accession number GSE189008. All relevant data and materials in this article are available from the corresponding authors upon reasonable request.

## Conflict of Interest

The authors declare that they have no competing interests.

## Acknowledgments

This work was partly supported by the Japan Society for the Promotion of Science KAKENHI grants (grant numbers 17J07299 to A.S., 19K16114 to A.S., 18H02427 to S.T., 17H06389 to C.F.); and the Japan Science and Technology Agency (JPMJER1902 to C.F.).

